# Oncomodulin Delays Age-Related Hearing Loss in C57 and CBA Mice

**DOI:** 10.1101/2021.06.11.448116

**Authors:** Leslie K. Climer, Aubrey J. Hornak, Kaitlin Murtha, Yang Yang, Andrew M. Cox, Preston L. Simpson, Andy Le, Dwayne D. Simmons

## Abstract

Ca^2+^ signaling is a major contributor to sensory hair cell function in the cochlea. Oncomodulin (OCM) is a Ca^2+^ binding protein preferentially expressed in outer hair cells of the cochlea and few other specialized cell types. Here, we expand on our previous reports and show that OCM prevents early progressive hearing loss in mice of two different genetic backgrounds: CBA/CaJ and C57Bl/6J. In both backgrounds, genetic disruption of *Ocm* leads to early progressive hearing loss as measured by auditory brainstem response (ABR) and distortion product otoacoustic emission (DPOAE). In both strains, loss of *Ocm* reduced hearing across lifetime (hearing span) by more than 50% relative to wild type (WT). Even though the two WT strains have very different hearing spans, OCM plays a considerable and similar role within their genetic environment to regulate hearing function. The accelerated ARHL of the *Ocm* KO illustrates the importance of Ca^2+^ signaling in maintaining hearing health.

## INTRODUCTION

Approximately 15% of American adults between the ages of 20 and 69 have high frequency hearing loss due to exposure to loud sounds, and 50% of Americans over 75 years old are affected by age-related hearing loss (ARHL), also known as presbycusis (https://www.nidcd.nih.gov/health/statistics). ARHL is a progressive loss of hearing sensitivity, impaired sound localization and decreased ability to understand speech, especially in noisy environments (Bielefeld et al. 2010, Wang and Puel 2020). Although progress has been made in defining some of the genetic and cellular functions that are disrupted by ARHL, little is known about its underlying causes. Sensory ARHL involves loss or degeneration of sensory hair cells and their neural connections in the cochlea (Altschuler et al. 2015, Jeng et al. 2020a, Jeng et al. 2020b, Lauer et al. 2012). Inner hair cells (IHCs) contact the majority of neural connections and transmit sound-induced impulses to the brain. Outer hair cells (OHCs) amplify and enhance IHC responses to sound. Recent studies in human temporal bones suggest that ARHL is largely affected by damage to inner ear sensory cells, particularly OHCs (Wu et al. 2020). During aging, as well as intense or prolonged noise exposure, damage to OHCs begins in basal, high frequency regions (Chen et al. 2009, Cruickshanks et al. 1998, Frisina 2009, Hu et al. 2006, Nelson and Hinojosa 2006, Wu et al. 2020). The loss of OHCs leads to elevated hearing thresholds and loss of cochlear frequency tuning.

Hair cell mechanotransduction, synaptic transmission and response to acoustic overstimulation depend on Ca^2+^ regulation (Fridberger et al. 1998, Jaramillo 1995, Lenzi and Roberts 1994). Outer hair cells regulate Ca^2+^ through an array of channels, transporters and mobile Ca^2+^buffers (Fettiplace and Nam 2019, Hackney et al. 2005). During development, OHCs show a dramatic shift in the expression profile of the mobile Ca^2+^ buffer family, EF-hand Ca^2+^ binding proteins (CaBPs). While certain CaBPs such as alpha-parvalbumin (αPV) are down regulated, oncomodulin (OCM), a member of the parvalbumin family, is upregulated in mouse OHCs soon after birth (P2-P3), supplanting other CaBPs as at the dominant Ca^2+^ binding protein (Climer et al. 2019, Henzl et al. 1997, Sakaguchi et al. 1998, Simmons et al. 2010, Thalmann et al. 1997, Yang et al. 2004). OCM is the only known mobile Ca^2+^ buffer that when deleted gives rise to a progressive hearing loss phenotype (Tong et al. 2016). In rodent OHCs, OCM expression is estimated to be in the millimolar range (2 - 4 mM), which is similar only to αPV expression found in fast twitch muscle (Hackney et al. 2005, Yang et al. 2004). The accelerated ARHL seen in *Ocm* KO mice strongly suggests that OCM is essential to the maintenance of hearing function in adult C57Bl/6J mice (Tong et al. 2016). Whether OCM has a role in ARHL is unknown.

Two mouse strains, the CBA/CaJ (CBA) mouse and C57Bl/6J (C57) mouse, are frequently used as models for late- and early-onset ARHL, respectively (Frisina et al. 2011, Greenwood 1990, Henry and Chole 1980, Henry and Lepkowski 1978, Jimenez et al. 1999, Li and Hultcrantz 1994, Spongr et al. 1997). The CBA mouse retains most of its hearing sensitivity up to 18 mo of age (in a 30-mo average lifespan), after which hearing declines progressively beginning in the high frequencies (Li and Borg 1991, Li and Hultcrantz 1994, Spongr et al. 1997). CBA mice do not show significant hair cell loss until around 18 mo with approximately 50% of OHCs remaining in the apex and base by 26 mo (Spongr et al. 1997). In contrast, the C57 mouse has been used as a model for accelerated (early-onset) ARHL. The C57 mouse demonstrates rapid onset and progression of hearing loss compared to the CBA mouse (Li and Borg 1991). The underlying pathology is progressive loss of cochlear sensory cells mediated by intrinsic apoptosis especially in basal hair cells after 6 mo with complete loss of basal OHCs by 12 mo (Francis et al. 2003, Li and Hultcrantz 1994, Someya et al. 2009, Spongr et al. 1997). Recently, Jeng and colleagues performed extensive analyses of the biophysical properties of aging inner and outer hair cells and their synapses in four genetic mouse strains with early or late ARHL onset (Jeng et al. 2021, Jeng et al. 2020a, Jeng et al. 2020b). They concluded that alterations in the mechanoelectrical transducer (MET) in the stereocilia bundle and OHC efferent synapses possibly contribute to the progression of ARHL, and that apoptosis of sensory cells may not be a contributing factor.

Given the unique expression profile of OCM in the cochlea, and the role that Ca^2+^ regulation may play in hearing function, we used the *Ocm* KO mouse model to explore the link between OCM and ARHL. The C57 strain is known to potentiate hearing loss of other Ca^2+^-regulatory gene mutations (Fettiplace and Nam 2019). Since C57 mice have an accelerated ARHL phenotype and the *Ocm* KO appeared to accelerate this phenotype even further, we hypothesized that the absence of OCM in mice on the CBA/CaJ genetic background will also induce an early progressive hearing loss phenotype but with a different time course. In this study, we compare age-related hearing decline in WT and KO mice of both backgrounds. In both mouse strains, the *Ocm* KO allele resulted in an early progressive hearing loss that was at least 50% of the normal hearing span, and a loss of efferent terminals and OHCs. Also in both strains, aged WT mice show alterations in the localization of OCM that coincide with the loss of OHCs and efferent terminals. This study suggests that OCM may indeed play a role in ARHL associated with OHC damage.

## MATERIALS AND METHODS

### Animals

All experiments were done in compliance with National Institutes of Health and institutional animal care guidelines and were approved by the Institutional Animal Care and Use Committee of Baylor University. *Ocm*^+/+^ (*Ocm* KO) mice were generated from spontaneous germline transmission of the KO allele from the parental line, *Actb^Cre^;Ocm^flox/flox^* on the C57Bl/6J background (Tong et al. 2016). To transfer the *Ocm* KO construct to the CBA/CaJ background, mice with a targeted deletion of *Ocm* were backcrossed onto the CBA/CaJ for 10 generations. At each generation, mice were genotyped for the *Cdh23-ahl* locus and only animals with the CBA/CaJ *Cdh23* allele were bred following procedures used by Liberman and colleagues (Liu et al. 2007). Confirmation of congenicity was done by whole genome scan (Jackson Laboratories, Bar Harbor, ME). Our C57 *Ocm* KO mice were crossed with *B6.129P2-Pvalbtm1Swal/J* on the C57Bl/6J background by Jackson Laboratories (Bar Harbor, ME). Homozygous CBA *Ocm* KO and C57 *Ocm* KO and wild-type (WT) littermates from each sex were used for all experiments.

### Antibodies

Tissues were incubated with primary antibody overnight at 37°C. Primary antibodies used: goat anti-oncomodulin (OCM, 1:500, Santa Cruz sc-7446), rabbit anti-prestin (1:5000, kindly provided by Robert Fettiplace), goat anti-choline acetyltransferase (ChAT, 1:500, Millipore, AB144P). All primary antibodies were labelled with species appropriate Alexa Fluor (ThermoFisher) or Northern Lights (R&D Systems) secondary antibody for 2 h at 37°C.

### Cochlear Function Assays

For measurement of auditory brainstem responses (ABRs) and distortion product otoacoustic emissions (DPOAEs), adult mice were anesthetized with xylazine (20 mg/kg, i.p.) and ketamine (100 mg/kg, *i.p*.). Acoustic stimuli were delivered using a custom acoustic assembly previously described by Maison and colleagues (2012). Briefly, two electrostatic earphones (EC-1, Tucker Davis Technologies) were used to generate primary tones and a Knowles miniature microphone (EK-3103) was used to record ear-canal sound pressure. Stimuli were generated digitally with 4s sampling. Ear-canal sound pressure and electrode voltage were amplified and digitally sampled at 20s for analysis of response amplitudes. Both outputs and inputs were processed with a digital I-O board (National Instruments PXI-4461). For measurement of ABRs, needle electrodes were inserted at vertex and pinna, with a ground electrode near the tail. ABR potentials were evoked with 5 ms tone pips (0.5 ms rise-fall with a cos2 onset, delivered at 35/s). The response was amplified (10,000), filtered (100 Hz-3 kHz), digitized, and averaged in a LabVIEW-driven data-acquisition system. Sound level was raised in 10 dB steps from 10 dB below threshold up to 80dB sound pressure level (SPL). At each sound level, 1024 responses were averaged (with stimulus polarity alternated), using an “artifact reject” whereby response waveforms were discarded when peak-to-peak amplitude exceeded 15V (e.g. electro-cardiogram or myogenic potentials). Threshold was defined as the lowest SPL level at which any wave could be detected, usually corresponding to the level step just below that at which the peak-to-peak response amplitude rose significantly above the noise floor. For amplitude versus level functions, the wave-I peak was identified by visual inspection at each sound level and the peak-to-peak amplitude computed. For measurement of DPOAEs at 2f1 - f2, the primary tones were set so that the frequency ratio, (f2/f1), was 1.2 and so that f2 level was 10 dB below f1 level. For each f2/f1 primary pair, levels were swept in 10 dB steps from 20 dB SPL to 80 dB SPL (for f2). At each level, both waveform and spectral averaging were used to increase the signal-to-noise ratio of the recorded ear-canal sound pressure, and the amplitude of the DPOAE at 2f1 - f2 was extracted from the averaged spectra, along with the noise floor at nearby points in the spectrum. Iso-response curves were interpolated from plots of DPOAE amplitude versus sound level. Threshold was defined as the f2 level required to produce a DPOAE at 0 dB SPL. We used a total of 25 female and 18 male WT and 13 female and 13 male KO CBA mice, and a total of 40 female and 38 male WT and 17 female and 25 male KO C57 mice. For both background strains, variances between female and male mice were negligible across the ages tested.

### Immunocytochemistry

For histological analysis and immunocytochemistry, anesthetized mice (Euthasol, 150 mg/kg, i.p.) were perfused transcardially with 4% paraformaldehyde (PFA) in 0.1M phosphate buffer (PBS) followed by PBS wash. After removal, cochleae scalae were flushed with PFA then rotated in fixative overnight at 4°C. Cochleae were decalcified in 0.1M EDTA for 3 - 5 days at 4°C with rotation. Cochleae were prepared as cochlear whole mounts of approximately 6 pieces of sensory epithelium or as 100 μm mid-modiolar sections embedded in gelatin-agarose solution (Maison et al. 2012, Simmons et al. 2010). Cochlear pieces were suspended in 30% sucrose for 30 - 60 minutes at room temperature with gentle shaking, frozen at −80°C for 30 min, then thawed for 30min at 37°C. Pieces were washed 3 times in PBS, then blocked in 5% normal horse serum (NHS) for 2 hours at room temperature. Images were acquired using the LSM800 microscope (Zeiss) using a high-resolution, oil-immersion objective (Plan-Apochromat 63x, 1.4 NA).

### Hair cell and efferent cluster counts

Low-power images of each microdissected piece of surface preparations were obtained with a 10x air objective (N.A. 0.3) on a LSM800 confocal microscope. Cytocochleograms were constructed from these images by tracing the cochlear spiral and superimposing hash marks using a custom ImageJ Measure Line plugin from Eaton-Peabody (https://www.masseyeandear.org/research/otolaryngology/eaton-peabody-laboratories/histology-core). The plugin superimposed frequency correlates on the microdissected spiral image by application of the cochlear frequency map for mice. Organ of Corti was imaged at the 8, 16 and 32kHz regions with the LSM800 confocal microscope using a 63x oil immersion objective (N.A. 1.4). At each of the desired locations, 3 adjacent microscopic fields (9-12 IHCs per field) were imaged with a 4-channel z-stack spanning the height of the hair cells to capture the stereocilia, nuclei and synaptic junctions. Hair cells were only counted if they had a nucleus or hair bundle/cuticular plate present. ChAT-labeled efferent clusters of terminal swellings were counted in equally spaced distance regions using a LSM5 confocal microscope using a 40x oil immersion objective (N.A. 1.2). In each cochlear location, we acquired two adjacent z-stacks (each spanning 112 μm of the cochlear spiral), taking care to span the entire region containing ChAT-positive terminals along the z-axis. Because of the difficulty counting ChAT-labeled terminals in older animals, we resorted to counts only of clusters of labeled efferent terminals. Counts were performed by a blinded observer.

### Statistical Analysis

All statistical analysis were performed in Prism (v9.1x GraphPad Software). All data were tested for homogeneity of variance. For multiple comparisons, Brown-Forsythe and Welch ANOVA test was applied when SDs are significantly different (p<0.05), followed by Dunnett’s multiple comparisons test. Mann-Whitney U-test was applied when normal distribution cannot be assumed. Mean values are quoted in text and figures as means ± SEM (DPOAE and ABR measurement) and means ± SD (missing OHCs counting). p<0.05 was selected as the criterion for statistical significance.

## RESULTS

### OCM prevents early progressive hearing loss

We previously showed that targeted deletion of *Ocm* in C57Bl/6j mice have elevated hearing thresholds by 3 - 4 mo (Tong et al. 2016). We wanted to compare the contribution of OCM to auditory function in mice with different genetic backgrounds, specifically, C57 mice with short hearing lifespans to CBA mice with longer hearing spans. We transferred the *Ocm* KO allele from the original C57 mouse line to the CBA background (*see methods*). To test hearing function, we compared ABR waveforms of C57 KO (**Figure 1A-D**) and CBA KO (**Figure 1E-H**) mice at early adult ages up to the ages where ABR potential responses were lost with age-matched WT C57 and CBA mice. ABR potentials at 8 and 12kHz frequencies are similar for WT and KO mice at 2 mo in both backgrounds (**Figure 1A-B, E-F**). *Ocm* KOs showed a lack of response under these same conditions as early as 4 mo in C57 mice (**Figure 1C-D)**and 7 mo in CBA mice (**Figure 1G-H**), whereas the WT animals demonstrated robust ABR potentials at 8 mo in both backgrounds. C57 *Ocm* KO ABR thresholds shifted 20 - 60 dB at middle and high frequencies between 1 and 3 mo of age (Tong et al. 2016). Tong et al (2016) showed a progressive elevation of ABR thresholds in the C57 KO. In CBA mice, we measured ABR thresholds at 3 different ages using 8, 16, and 32kHz (**Figure 1I-J**). WT and KO animals showed similar thresholds at 2 mo, but KO animals showed threshold shifts by 5 mo, particularly at higher frequencies, and were essentially deaf by 7 mo (**Figure 1I**). In the CBA KO, 2 - 7 mo DPOAE thresholds were significantly different from each other at 16kHz (Welch’s ANOVA: p = 0.0003; Dunnett’s post-test: p = 0.0002 for 2 −7 mo). In contrast, WT animals demonstrate similar ABR thresholds at low and middle frequencies from 2 - 7 mo, and only show substantial threshold shifts in the high frequencies at 5 - 7 mo (**Figure 1J**). At 16kHz in the CBA WT, there was no significant difference in ABR thresholds up to 7 mo (Welch’s ANOVA: p = 0.104, followed by Dunnett’s test).

**Figure 1.**
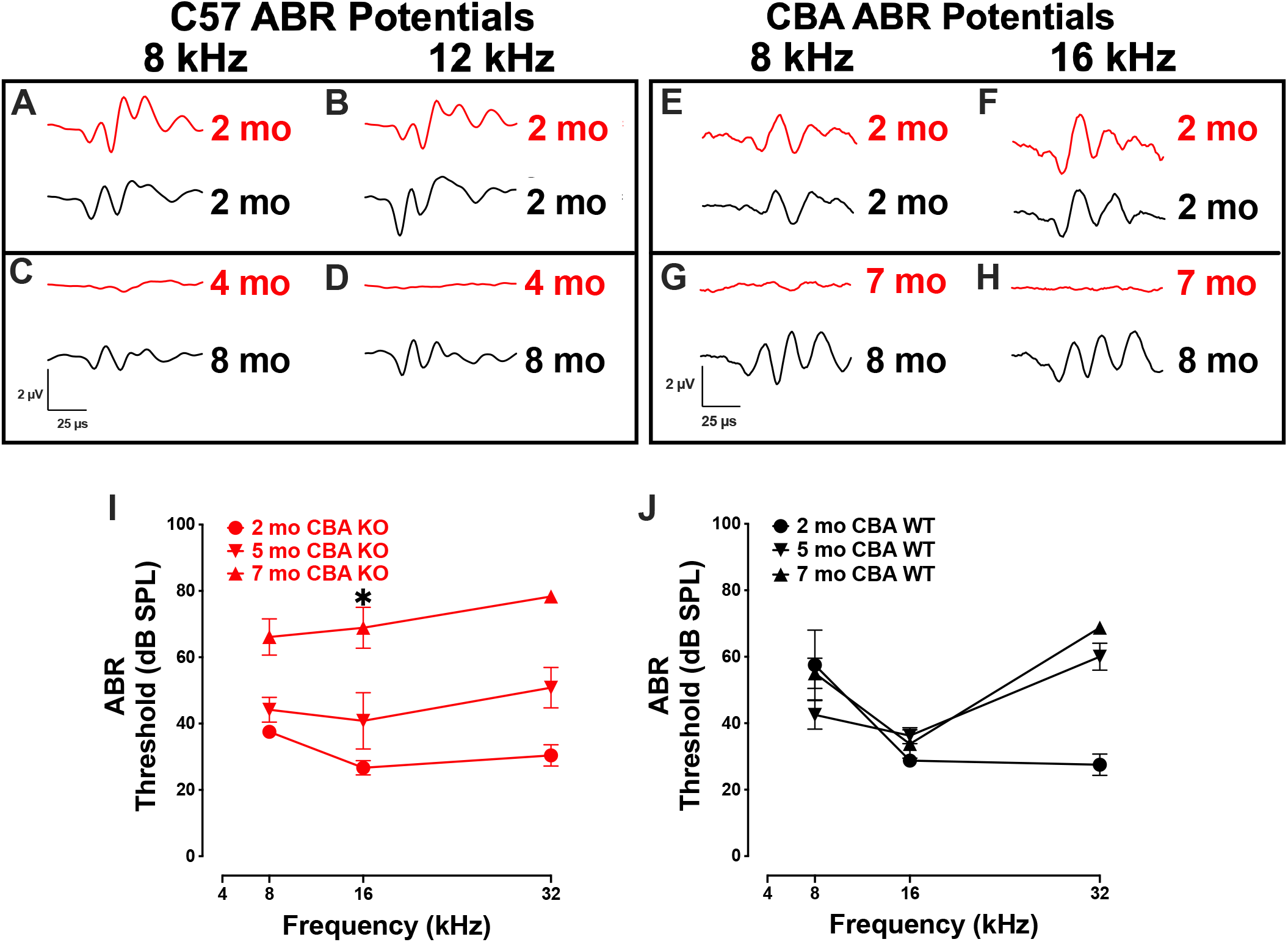
Loss of OCM alters ABR potentials and thresholds. A-D) Representative ABR waveforms from WT (black) and KO (red) C57 mice produced using 8 and 12 kHz tone bursts given at 80 dB SPL. Waveforms for 2 mo and 8 mo C57 WT, and 2mo and 4mo C57 KO animals. E-H) Representative ABR waveforms from WT (black) and KO (red) CBA mice produced using 8 and 16 kHz tone bursts given at 80 dB SPL. Waveforms for 2 mo and 8 mo CBA WT, and 2 mo and 7 mo CBA KO animals. I-J) Mean ABR thresholds for 2 - 7 mo CBA KO (I, *n* = 6-9 per age) and WT mice (J, *n* = 4 per age). Asterisks represent significance (p<0.05) from 2 mo at 16 kHz.

To confirm that germ-line transmission of the *Ocm* KO in C57Bl/6J mice did not change the original hearing loss phenotype described by Tong et al (2016), we measured DPOAE thresholds in age-matched KO and WT C57 and CBA mice over the course of their hearing span (**Figure 2A-F**). The C57 WT control mice showed little threshold shifts between 1 and 5 mo. The C57 KO mice show a progressive elevation of DPOAE thresholds from 1 to 6 mo (**Figure 2A**). At 16kHz, there were significant differences in DPOAE thresholds between 1 - 3 mo and 3 - 6 mo (Welch’s ANOVA: p < 0.0001; Dunnett’s test: p = 0.009 for 1 - 3 mo, p= 0.002 for 3 - 6 mo). In the C57 KO between 3 and 6 mo, we observed sizeable threshold shifts in all but the lowest frequencies (8 - 32 kHz), while in the WT, sizeable threshold shifts were observed between 5 and 16 mo (**Figure 2A, B)**. The largest threshold shifts in the C57 KO were in the middle frequencies. Although C57 KO mice were at or near the limit of DPOAE threshold detection by 6 mo, the C57 WT mice had measurable DPOAEs in most frequencies at 16 mo. In the C57 WT, there were no significant differences in 16kHz thresholds of 1,4 and 5 mo mice (Dunnett’s test). There was a significant difference in thresholds between 1 mo and 16 mo at 16 kHz (p = 0.009, Dunnett’s test). Since OCM is predominantly expressed in OHC of the organ of Corti at these ages, these results confirm that OHC defects are primarily responsible for the differences in ABR potentials between the *Ocm* KO early onset hearing loss and the WT.

**Figure 2.**
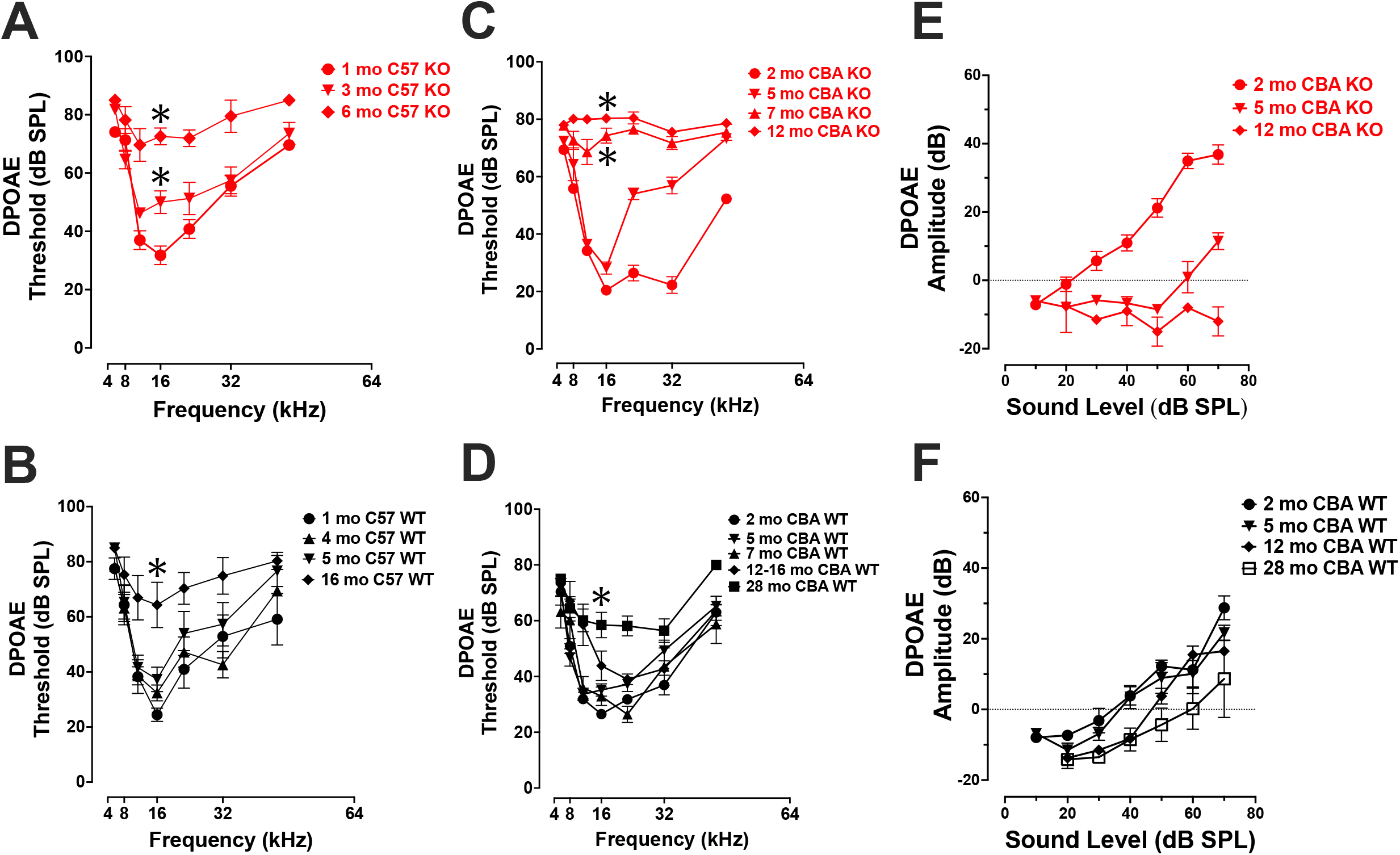
*Ocm* KO mice show progressive elevation in DPOAE thresholds. A-B) Mean ± SEM DPOAE thresholds as a function of *f*2 frequency for 1 - 6 mo C57 KO (A, *n* = 23) and 1 - 16 mo WT mice (B, *n* = 33). C-D) Mean ± SEM DPOAE thresholds as a function of *f2* frequency for CBA mice. C) DPOAE thresholds from 2 - 12 mo CBA KO (*n* =31) animals and D) 2 - 28 mo WT (*n* = 49) animals. E-F) Mean ± SEM DPOAE input/output functions from E) KO and F) WT animals. Asterisks represent significance (p<0.05) from 1 - 2 mo at 16 kHz.

Similar to their ABR threshold responses (**Figure 1I**), CBA KO animals showed middle- to high-frequency DPOAE threshold shifts from 2 to 5 mo (**Figure 2C**). The KOs also showed nearly maximum threshold shifts at all frequencies from 7 - 12 mo. At 16 kHz, DPOAE thresholds were not statistically different between 2 and 5 mo but were significantly different between 2 and 7 mo and 2 - 12 mo (Welch’s ANOVA: p < 0.0001; Dunnett’s post-test: p < 0.0001 for 2 - 7 mo, p < 0.0001 for 2 – 12 mo). In contrast, WT mice maintained healthy threshold levels at least up to 16 mo of age and still produced measurable DPOAE threshold responses in most frequencies by 28 mo (**Figure 2D**).

At 16 kHz in CBA mice, only DPOAE thresholds between 2 and 28 mo were significantly different (p < 0.0005, Dunnett’s test). Interestingly, young adult KO mice (2 mo) show enhanced DPOAE input-output functions compared with WT animals suggesting enhanced electromotility at least for higher frequencies. We compared DPOAE input-output functions at 32 kHz across ages (**Figure 2E-F**). The KO animals required less signal (dB SPL) to produce a DPOAE output that crosses threshold (**Figure 2E**). This efficiency is lost by 5 mo, and by 12 mo the KO input-output responses do not cross threshold. WT animals lose their input-output efficiency between 5 and 12 mo, requiring higher dB SPL input to cross threshold than at younger ages (**Figure 2F**).

Taken together, *Ocm* KO reduces the hearing lifetime (hearing span) of mice regardless of genetic background (**Figure 3**). Though CBA mice are a well-established mouse model that hears throughout most its life, *Ocm* deletion in OHCs reduces the hearing span to less than half of the time of WT counterparts with severe hearing dysfunction in the first 1/3 of life (**Figure 3, 2nd bar**). Deletion of *Ocm* similarly reduces the hearing span of C57 mice, but at an earlier age. When comparing hearing thresholds between the two genetic backgrounds, loss of OCM in the CBA background recapitulates a similar hearing phenotype of C57 background (**Figure 3, middle bars**). These data demonstrate that OCM is an essential contributor to hearing health by delaying progressive hearing loss as a function of age.

**Figure 3.**
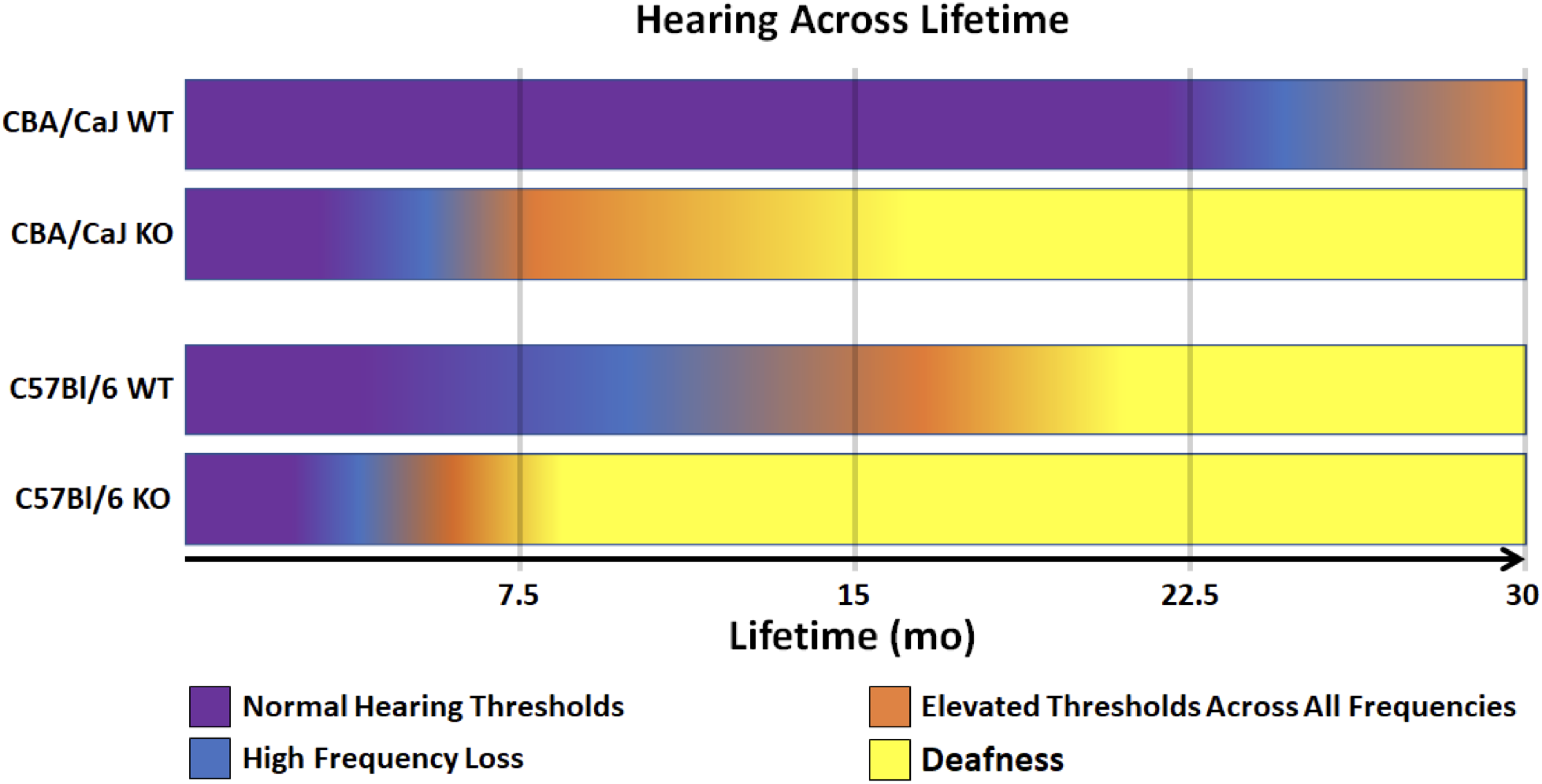
Model of WT and KO hearing spans in CBA and C57 backgrounds. Comparison of changes in hearing outputs after OCM loss across a 30-mo average lifetime. CBA WT mice show high frequency loss after 20 mo (blue). CBA KO mice show high frequency loss in less than half the time (5 - 7 mo) which is earlier than when the C57 WT mice lose high frequencies (7 - 10 mo). C57 KO mice lose high frequencies at 3 - 4 mo. Deafness (yellow) is defined as maximum elevation of DPOAE thresholds at all frequencies tested (5.6 - 45.2 kHz range).

### OCM and prestin are expressed in defective OHCs

Since *Ocm* KO mice appear to accelerate early progressive hearing loss, we wanted to compare aging OHC phenotypes in the WT and KO mice across genetic backgrounds. In CBA animals, WT animals manifest substantial OHC loss after 1 year (**Figure 4A-C, I**). Interestingly, OCM expression is maintained even when there is substantial OHC loss. However, the subcellular localization and intensity of OCM immunostaining varies with age and frequency region across genetic background. In CBA WT mice, we find OCM immunoreactivity is qualitatively more intense in the nucleus than in the cytosol of 2-month-old animals in the 8, 16 and 32kHz regions (**Figure 4A-C**). OCM expression is more evenly divided between the nucleus and cytosol by 12 mo, but there are a few cells with intensely labeled OCM in the cytosol and lateral membrane (**Figure 4A-C**, **12mo^*^**). This phenomenon is more extreme at 30 mo with the majority of the remaining OHCs showing intense cytosolic OCM labeling particularly in the 16 and 32kHz regions (**Figure 4B-C**, 30mo). Additionally, the WT OHC damage and/or loss was more pronounced in apical regions (**Figures 4 and 5**). High resolution images of 3 mo and 22 mo WT animals demonstrate the intense cytosolic immunolabeling of OCM at older ages (**Figure 4D, green**). Since we know that WT animals have poorer hearing thresholds from 12 - 30 mo, it is possible that the OCM expression profile is a feature of dysfunctional/aging OHCs. Older animals still maintain expression of the OHC specific motor protein prestin, responsible for mechanotransduction, though this too may be dysfunctional (**Figure 4D, white**). WT animals maintain robust OHC numbers throughout the first year of life and are only missing approximately 50% of the OHCs along the entire cochlear spiral by 30 mo (**Figure 4I, top**). By contrast, KO animals have lost the majority of OHCs by 12 mo (**Figure 4I, top**). Similar to our previous report (Tong et al. 2016), WT OHC numbers were maintained through 5 mo of age in the C57 background (**Figure 4F-I**) unlike *Ocm* KO animals (**Figure 4H, I bottom**). Prestin was also abundant in the remaining OHCs of aged WT mice (**Figure 4F-G**) and KO OHCs at 5 - 6 mo (**Figure 4D, H**). Similar to the CBA WTs, OCM is expressed more intensely in the remaining C57 OHCs with age (**Figure 4F**) and across all frequencies (data not shown).

**Figure 4.**
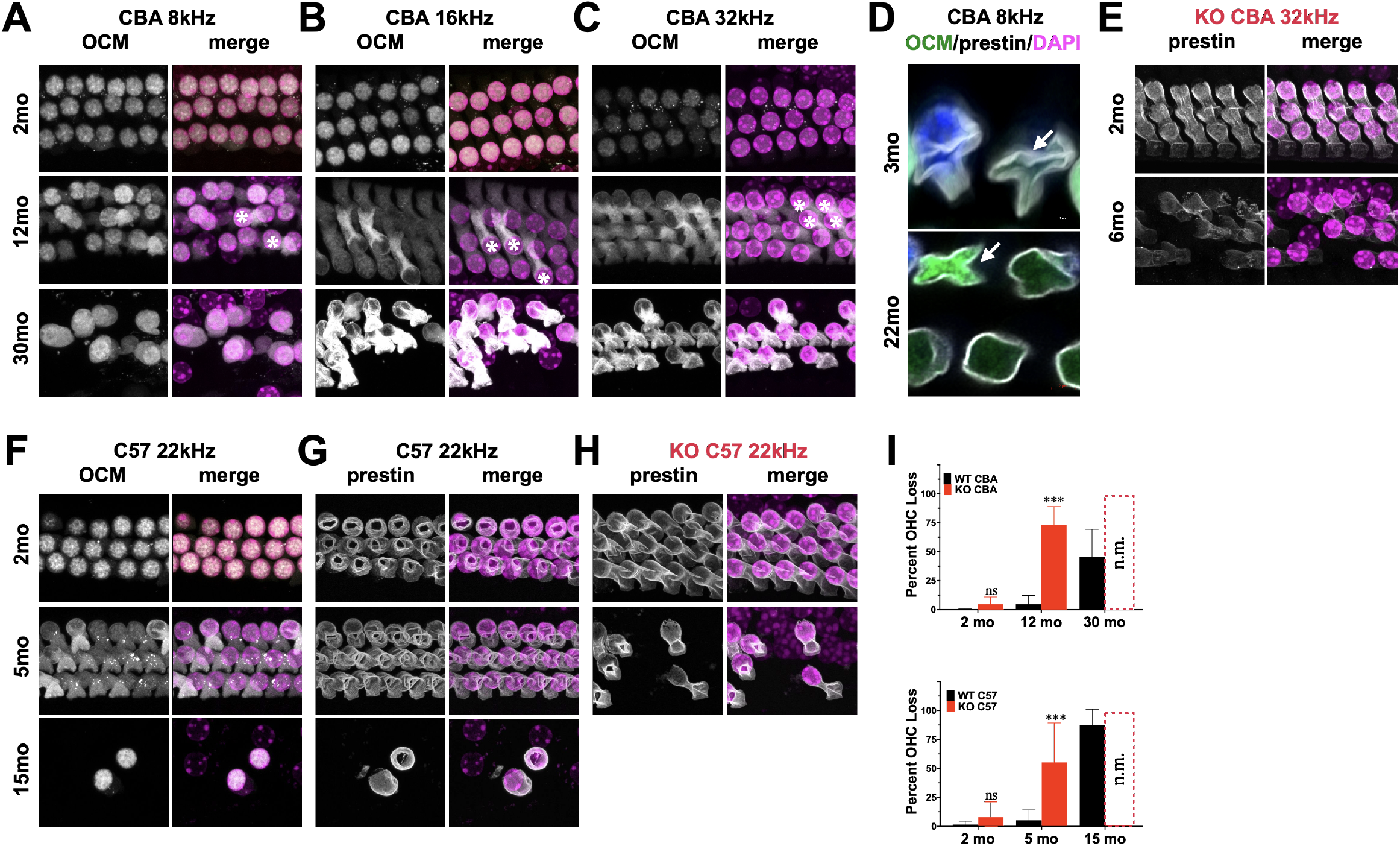
OCM and Prestin are expressed in aged OHCs. A-C) Surface preparations from 2 mo, 12 mo and 30 mo CBA WT mice stained for OCM (gray) in the A) 8kHz, B) 16kHz and C) 32 kHz regions. Asterisk represents enhanced OCM immunoreactivity. D) Upper panel: 3 mo-old CBA/CaJ WT OHC at 8 kHz. Arrow indicates collapsed OHC with low OCM expression. Lower panel: 22 mo-old CBA/CaJ OHC at 32 kHz. Collapsed OHC with saturated OCM expression (arrow). Remaining OHCs have very low intensity OCM expression. E) 32 kHz region surface preparations from CBA KO mice stained for prestin at 2 mo and 6 mo. F-G) 22 kHz region surface preparations from 2 mo, 5 mo and 15 mo C57 WT mice stained for F) OCM and G) prestin. H) 22 kHz region surface preparations for C57 KO mice stained for prestin at 2 mo and 5 mo. I) Upper panel: CBA WT and KO OHC loss quantified from surface preparations. Lower panel: C57 WT and KO OHC loss quantified from surface preparations. OHC loss was not measured from the KOs at 30 mo and 15 mo (n.m.). Loss was averaged from samples taken along the entire cochlear spiral. Error bars are S.D. of 3 samplings. Triple asterisks represent significance (p<0.05) from aged-matched WT. n.s. is not significant.

**Figure 5.**
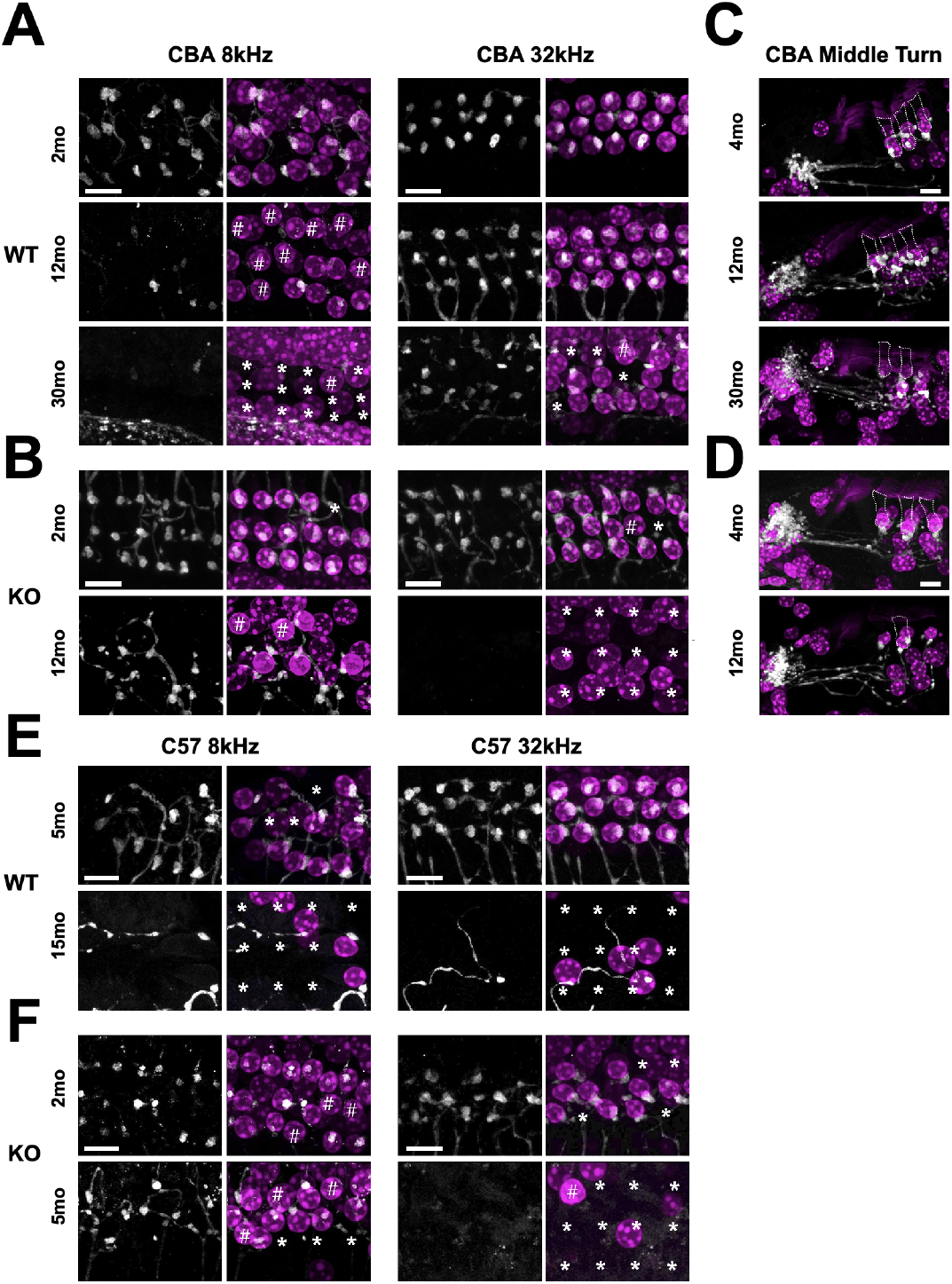
Efferent clusters are lost at the extreme ends in aging WT and KO mice. Surface preparations showing ChAT-labeled efferent clusters in the 8 and 32 kHz regions of A) CBA WT and B) CBA KO mice. ^*^ Missing OHCs, # OHCs without ChAT clusters. C-D) Mid-modiolar sections of ChAT-labeled clusters in the middle turn of C) CBA WT and D) CBA KO mice. Dotted lines represent OHC borders from adjacent OHC in each row. Scale bars represent 10 μm.

### Efferent connections are lost with age and in WT and KO animals

Synaptic dysfunction, rewiring and OHC loss are common phenotypes of the aging cochlea. Activation of cholinergic medial olivocochlear (MOC) efferent axons protects OHCs from excessive noise and Ca^2+^ by modulating OHCs directly and altering their Ca^2+^-sensitive motility (Boero et al., Frolenkov 2006, Maison et al. 2003, Simmons 2002). Loss of MOC efferents could enhance the dysfunctional state of aging OHCs and thus result in increased hearing thresholds (Fu et al. 2010). Therefore, we looked for the presence and absence of ChAT-labeled efferent terminal clusters in aging WT and KO animals. In both CBA and C57 mice, we co-labeled surface preparations (**Figure 5A-B, E-F**) and mid-modiolar cochlear sections (**Figure 5C-D, CBA only**) with DAPI to stain the nuclei and ChAT to label efferent fibers and terminal clusters. In CBA WT and KO mice, ChAT-labeled clusters were abundant at 2-4 mo. However, in older WT and KO animals, we observed many instances of OHC loss and absence of ChAT labeling on remaining OHCs. Apical OHCs lose ChAT clusters by 12 mo in WT animals (**Figure 5A#**). At 30 mo in the 8kHz region, the few remaining OHCs do not have ChAT-labeled clusters (**Figure 5#**). The KOs show some hair cell loss and efferent loss as early as 2 mo in the 32kHz region, but the most dramatic efferent loss occurs in both apical and basal regions by 12 mo (**Figure 5B^*^#)**. Both KO OHC and efferent loss appear similar to recent data collected from young and old human temporal bones (Wu et al. 2019, Wu et al. 2020). The WT OHC damage and/or loss was more pronounced in apical regions while the KO OHC loss was pronounced in basal regions.

In C57 mice, the pattern of OHC and efferent loss was more pronounced. The WT apical OHC and efferent loss was evident as early as 5 mo and by 15 mo both apical and basal OHCs and efferent clusters were mostly absent (**Figure 5E**). In the KO at 2 mo, we observe apical efferent loss and basal OHC loss (**Figure 5F**). At 5 mo, the majority of basal OHCs were absent and there were also no efferents clusters. Apical regions had less OHC and efferent loss. WT mice showed similar apical and basal OHC and efferent loss while the KO showed more OHC and efferent loss in basal regions.

### OHC dysfunction occurs prior to cell death

We analyzed the loss of OHCs and ChAT-labeled efferent connections in relation to specific DPOAE frequency regions in CBA KO animals (**Figure 6**). We co-labeled surface preparations with antibodies to prestin and ChAT (**Figure 6A-F**). In young animals, we found robust DPOAE responses at 16 and 22kHz (**Figure 6H, purple**) with the majority of hair cells present except in the extreme base (**Figure 6G, purple**). By 6 - 7 mo, responses were near the measurement ceiling at these frequency regions (**Figure 6H, green**) and yet there are still hair cells present, particularly at 16kHz regions (**Figure 6G, green**). Indeed, DPOAE responses were elevated near maximal sound levels across the entire cochlea at 7 mo, but the majority of hair cells in all three OHC rows are present in the 4-16kHz regions and prestin was present in the remaining cells. ChAT was also present in the majority of the remaining cells, though not all (**Figure 6C-D#**). Efferent loss was progressive, with the greatest loss occurring by 7 mo in the middle and high frequency regions (**Figure 6I**). We plotted the percentage of efferent bouton clusters present (2 and 7 mo comparison) and the percent OHC present as a function of cochlear frequency (**Figure 6J**). Middle frequency regions of the cochlea show the least efferent loss. The greatest loss of efferent clusters occurs in the regions of greatest OHC loss. Apical regions of the cochlea also show a loss of efferent clusters with little OHC loss. Taken together, there is a complex relationship between OHC loss, efferent terminal loss and hearing loss across cochlear frequency regions. In agreement with previous literature, high frequency regions are the most susceptible to trauma.

**Figure 6.**
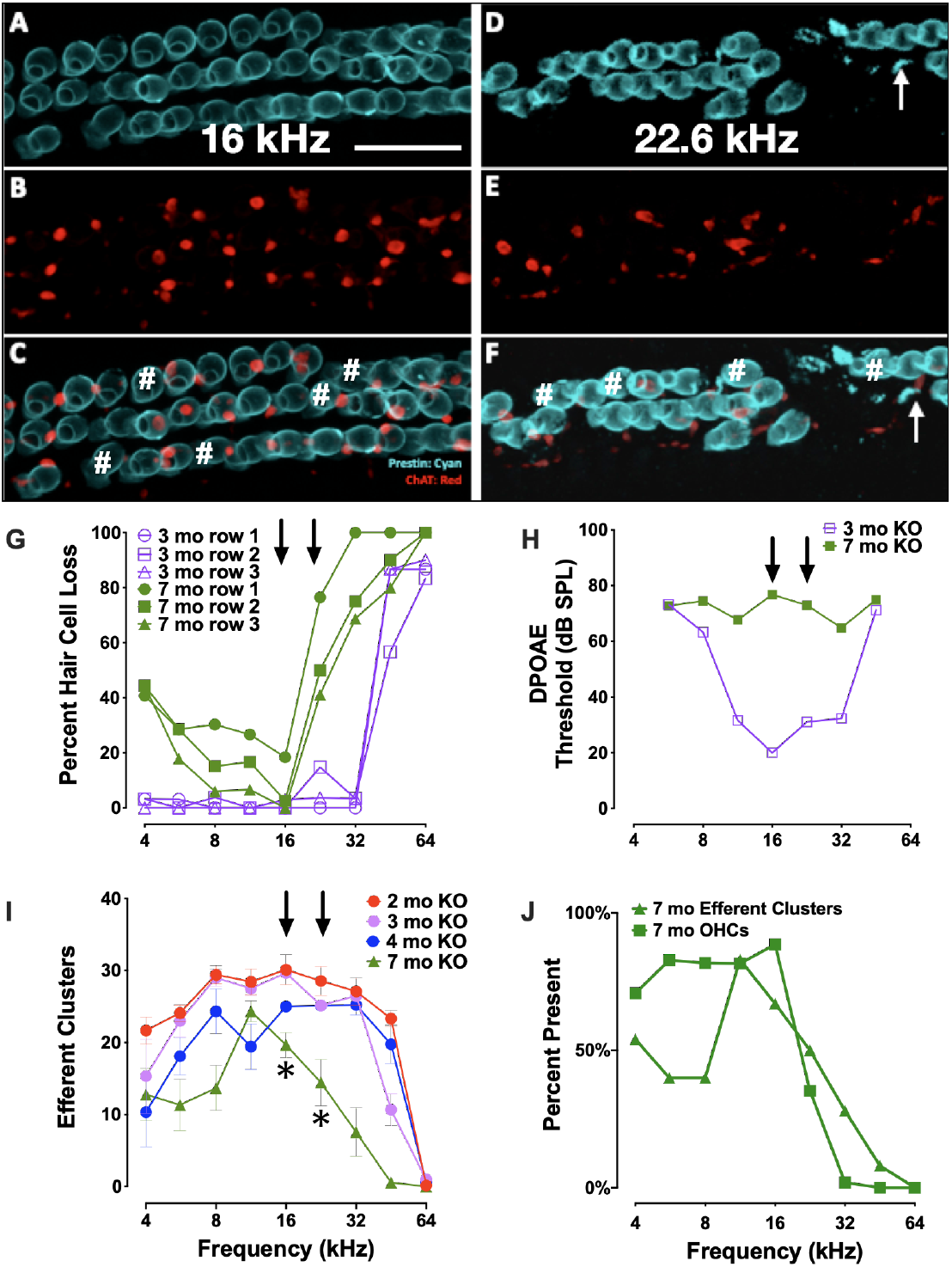
Elevated thresholds occur even when OHCs and efferent clusters are present. A-F) Surface preparations of 7 mo CBA KO cochlea co-labeled with prestin (teal) and ChAT (red) in the 16 and 22 kHz regions of CBA KO mice. # OHCs without ChAT clusters. Arrows point to OHC fragments. G) % of OHCs missing from each OHC row at 3 mo and 7 mo. Arrows point to 16 and 22 kHz frequencies. H) DPOAE thresholds of animals in (G). Arrows point to 16 and 22 kHz frequencies. I) Quantification of the number of ChAT-labeled efferent clusters from CBA KO animals from 2 - 7 mo (n = 3/age). J) Comparison of efferent and OHC loss in 7 mo KO mice (n = 3). Scale represents 40 μm. Asterisks represent significance (p<0.05) from 2 mo at 16 and 22 kHz.

## DISCUSSION

In this study, we assessed the role of OCM, an EF-hand CaBP predominately expressed in OHCs, in ARHL using two different genetic strains of mice: CBA/CaJ and C57Bl/6J. The CBA mouse has a substantially later onset of ARHL compared to the C57 mouse. First, we engineered an *Ocm* KO mouse using C57Bl/6J and then backcrossed the *Ocm* KO onto CBA/CaJ. Compared to their WT controls, both *Ocm* KO strains demonstrated an earlier progressive elevation of hearing thresholds, loss of efferent synaptic contacts, and OHC loss. Our results in the *Ocm* KO suggest that alterations in Ca^2+^ signaling lead to OHC dysfunction before loss of efferent synaptic contacts and/or OHC loss. Although we do not have direct evidence, our results suggest that changes in the presence of efferent terminals may occur prior to OHC loss. The WT and *Ocm* KO mice share common aging characteristics despite differences in their temporal progression of hearing loss.**Figure 7**represents a simple model illustrating the general structural similarities between aged WT and young *Ocm* KO animals across frequencies. Both aged WT and young KO show OHCs and ChAT labeled efferent connections are lost with age. Importantly, even when ABR and DPOAE thresholds are highly elevated or not measurable across frequencies, the majority of OHCs and efferent terminals are still present in most frequency regions in both aged WT and younger *Ocm* KO animals, suggesting compromised OHC electromotility function. At least in OHCs, it is possible that disturbances in Ca^2+^ signaling caused by the removal of OCM lead to the early activation of a general aging process. The absence of OCM in OHCs appears to accelerate ARHL and mimics human presbycusis, which typically displays loss of nerve fibers and hair cells at apex and base (Wu et al. 2019). Future studies assessing both IHC and OHC survival and their neural connections in *Ocm* KO animals will be essential to identifying other characteristics of human presbycusis.

**Figure 7.**
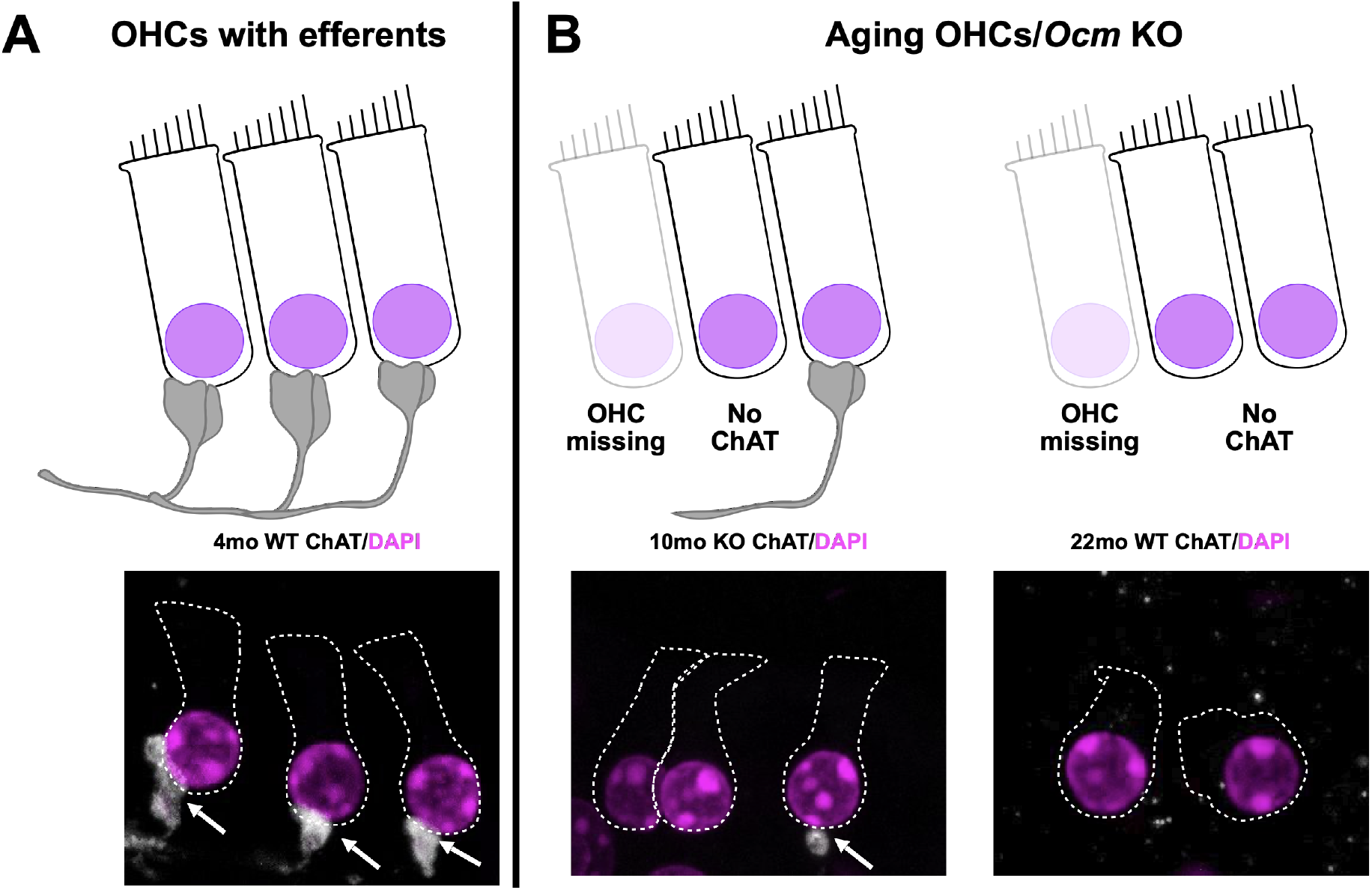
Model of common *Ocm* KO and aging characteristics. A) Young adult CBA WT and KO animals have organized OHC rows with efferent clusters present on all cells. B) During aging, OHCs are lost and not all of the remaining cells will possess efferent synaptic connections. In the WT animals, this occurs after 1 year, but the KOs show efferent and OHC loss much earlier. This demonstrates that dysregulated Ca^2+^ signaling can enhance the effects of aging.

### Our work extends findings of Tong et al. (2016)

Oncomodulin plays a major role in the maintenance of OHC intracellular Ca^2+^ levels. Outer hair cells have very high levels of OCM with estimates reaching as high as 2-4 mM suggesting that OCM provides the bulk of Ca^2+^ buffering in OHCs (Hackney et al.). Ca^2+^ may be involved in regulating the motor capability underlying OHC cochlear amplification. The high concentration of OCM in OHCs, similar only to αPV in skeletal muscle, may protect against deleterious consequences of Ca^2+^ loading after acoustic overstimulation. Thus, without OCM to buffer the Ca^2+^ levels in OHCs, cytoplasmic Ca^2+^regulation could be severely disrupted and lead to OHC dysfunction and loss if nothing effectively compensates for its absence (Climer et al. 2019). To date, unlike previous genetic disruption studies of other mobile Ca^2+^ buffers that show little, if any, impact on hearing, OCM is the only known CaBP for which targeted deletion causes progressive hearing loss (Tong et al. 2016), suggesting the essential role of Ca^2+^ signaling in maintaining hearing health. Within the inner ear, targeted deletion of any of the other major EF-hand CaBPs (e.g., calbindin D28k, calretinin and αPV) show no hearing loss phenotype (Airaksinen et al. 2000, Schwaller et al. 2002). Using a targeted deletion of *Ocm*, Tong et al (2016) demonstrated that the lack of OCM led to progressive cochlear dysfunction beginning after 1 mo and resulted in ABR threshold shifts and loss of DPOAEs by 4 - 5 mo of age. Functionally, the absence of OCM mimicked an accelerated aging or noise damage process. The presence of normal ABR and DPOAE thresholds in the *Ocm* KO at 1 mo suggested that OCM was not essential for the development of cochlear function, but critically protects OHCs from damage in the adult ear. Similarly in the present study, we used a C57 mouse with germ-line deletion of OCM. The majority of C57 KO mice had significantly higher ABR and DPOAE threshold responses at 5 - 7 mo across all frequencies, but the absence of responses could occur as early as 4 mo at some frequencies. Compared to the WT mice at 5 mo, these *Ocm* KO mice also exhibited increased OHC loss especially in basal regions and altered OHC prestin immunoreactivity. In fact, the *Ocm* KO mice had hearing thresholds, OHC loss and prestin immunoreactivity comparable to 15 mo old WT C57 mice.

### Comparison of Ocm KO on CBA with C57

The two most utilized mouse models for studies of ARHL are the C57Bl/6J and CBA/CaJ strains. The C57 strain has a rapid, high-frequency hearing loss and loses most of its high-frequency hearing during the first 12 mo. In contrast, the CBA strain loses its hearing slowly with age. At early adult ages from 1 - 2 mo, functional measures of hearing for these two mouse strains are virtually indistinguishable. In the present study, the CBA *Ocm* mutant mice present with an early onset ARHL as seen in the C57 WT and KO mice. Our CBA *Ocm* mutants, beginning around 5 - 7 mo, and aged WT mice show similar hallmarks of ARHL: 1) progressive elevation of DPOAE thresholds, 2) loss of OHC efferent terminals, and 3) OHC loss. In *Ocm* KOs on a CBA background, our data show a progressive elevation of DPOAE thresholds over 12 mo. In contrast, WT hearing thresholds and response magnitudes change little from 2 - 12 mo. Similar to our observations in the C57 strain, young (2 mo) CBA *Ocm* KO mice had DPOAE thresholds nearly identical to WT controls at lower frequencies and somewhat enhanced at higher frequencies. In older CBA *Ocm* KO mice, maximum DPOAE threshold shifts occurred at frequencies 16 kHz and higher, which is within the region of greatest hearing sensitivity in these animals. *OCM* KO mice had significantly higher thresholds at all measured frequencies between 5 - 7 mo. At 12 mo, DPOAE thresholds were only slightly elevated in WT control mice and were not measurable in the *OCM* KO. In the present study, WT CBA mice had measurable DPOAEs up to 28 mo and we did not find significant ARHL until after 28 mo. These results suggest that *OCM* deletion in CBA mice leads to an accelerated ARHL phenotype that is faster than the WT C57, but more delayed than in C57 KO mice. Although the specific temporal pattern of hearing loss differs, the results were similar to our previous report and validate that OCM expression is critical for either the maintenance of cochlear function and/or protecting OHCs from damage as previously hypothesized (Tong et al. 2016). Further, having a similar early progression of hearing loss across genetic strains argues that OCM possibly mediates sensitivity to ARHL. As illustrated in **Figure 3**, comparing the effect of OCM deletion on hearing thresholds in CBA and C57 mice reduces the period of measurable hearing thresholds by over 50%.

### Comparison of Ocm KO with ARHL

In *Ocm* KO mice with high DPOAE thresholds, the majority of OHCs are still present. Although dysregulated Ca^2+^ signaling is a known contributor to apoptotic cell death, OHC loss occurs well after the loss of DPOAEs in the *Ocm* KO mouse. Since DPOAE responses are a direct measure of OHC function, elevated DPOAE thresholds suggest the remaining OHCs in the *Ocm* KO must be either “silent” or unresponsive to sounds. In CBA mutants, supra threshold DPOAE responses show rapid attenuation between 2 and 5 mo. At 32 kHz, 2 mo sensitivity to sounds deteriorated nearly 40 dB by 5 mo even though the majority of OHCs were still present. The lack of DPOAEs might suggest a decrease in expression of the motor protein prestin, as loss of prestin is correlated with elevated DPs (Liberman et al. 2002). However, this is unlikely due to the robust prestin immunoreactivity in these unresponsive OHCs. The lack of DPOAEs but presence of prestin protein would suggest that Ca^2+^ dysregulation makes OHCs dysfunctional in other ways such as in their biophysical characteristics. Recent studies of aged OHCs are consistent with the idea that OHCs undergo a period of dysfunction prior to OHC loss. In a study of aged OHCs from early onset and late onset ARHL mouse models, Jeng and colleagues suggested that age-related OHC dysfunction is not due to apoptosis (Jeng et al. 2021). They primarily focused their studies on OHCs from the 9-12kHz region. At 12 - 13 mo irrespective of whether OHCs were from early and late ARHL mice, they found OHCs had similar biophysical properties related to their peak current-voltage relationships, potassium currents, membrane capacitance, and membrane voltage measurements. With the exception of membrane voltage, they showed that these biophysical properties decreased similarly with age, being highest in the youngest animals. In addition to OHC loss, they found decreases in OHC size as indicated by decreases in membrane capacitance, OHC ribbons, and the mRNA expression of *Slc26a5* and *OCM* (Jeng et al. 2021). Most, if not all, of these features changed similarly, independent of the ARHL onset suggesting that they are general features of aging in OHCs, and not necessarily related to the OHC dysfunction observed in ARHL. Thus, it is possible that OHC-dependent ARHL is associated with disruption of cellular processes of transduction or motility. In the present study, we observed both prestin and OCM immunoreactivity in aged WT C57 and CBA OHCs (**Figure 4**). However, both proteins label more intensely with age (**Figure 4**), most likely either due to changes in cellular volume or expression. The seemingly enhanced expression of OCM could simply represent an artifact of shrunken OHC volume or it might convey that there is increased Ca^2+^ activity (**Figure 4D**). In this regard, altered Ca^2+^ buffering could change the dynamics of organelles (e.g., mitochondria and endoplasmic reticulum), which store and release Ca^2+^ as well as alter ATP utilization. Although it is possible the overall mRNA expression of *Ocm* and *Slc26a5* (prestin) in the cochlea decreases with age as reported by Jeng et al (2020b), the ramifications for protein levels remain unclear. A more relevant question is whether their functions are somehow compromised. Jeng et al (2020b) measured non-linear capacitance in aged OHCs and found that it decreased, but when normalized, the decrease appears independent from an ARHL phenotype. It is possible that smaller cell volumes due to an age-related shrinkage causes a relative increase in prestin protein levels. A tangential question is raised: though prestin is still present in aged OHCs, how functional is it? Overexpression of dysfunctional prestin in the mutant knock-in is more deadly to OHCs than complete lack of prestin (Keller et al. 2014). Our finding that OCM localization appears to undergo changes associated with age in WT mice might indicate age-associated changes in Ca^2+^ activity. It seems one of our challenges will be separating general features of aging in OHCs from those that lead to ARHL.

### OHC efferent innervation during ARHL

Sound- or electrically-induced activity of the cholinergic MOC pathway suppresses OHC responses leading to reduced cochlear output and protection from acoustic injury (Maison and Liberman 2000, Rajan and Johnstone 1989, Zheng et al. 1997) Previous studies have shown that the function of the MOC system declines with age prior to OHC degeneration in both humans and mice (Boero et al. 2020, Jacobson et al. 2003, Kim et al. 2002, Sun and Kim 1999, Zhu et al. 2007). In adult mice, OHCs are primarily innervated by the cholinergic MOC neurons, which modulate amplification of the cochlear partition (Guinan 2006). At MOC - OHC synapses, release of acetylcholine (ACh) causes Ca^2+^ entry through α9α10 nicotinic ACh receptors (α9α10 nAChRs). The Ca^2+^ influx activates a hyperpolarizing current through Ca^2+^-activated small conductance K^+^ (SK2) channels on the OHC and causes OHCs to elongate. The overall effect of OHC elongation is believed to be protective. In both our *OCM* KO and in aged WT controls, the presence of cholinergic efferent terminals contacting OHCs decreased in both low and high frequency regions with age (**Figure 5**). Jeng et al (2021) also investigated whether the efferent innervation was retained in the aged apical cochlea of early and late onset ARHL mice. Although Jeng et al (2020b) found no significant change in either the percentage of OHCs with SK2 immunoreactivity or the percentage of SK2 puncta juxtaposed to cholinergic terminals, they did note that some apical OHCs showed only efferent terminals or SK2 puncta. It has been known for some time that a reduction in OAEs correlates with impaired cochlear responses (de la Cruz et al. 1998, Zhu et al. 2007). If efferent terminals or SK2 puncta are lost, OHCs would lose that protection and have increased susceptibility to noise damage. MOC-mediated resistance to ARHL has been induced by enhancing α9α10 nAChR complexes on OHCs (Boero et al. 2020).Thus, the efferent loss or decline during aging could exacerbate ARHL. Future studies should investigate the time course of efferent synaptic loss on the pre- and post-synaptic OHC junction.

## CONCLUSION

By comparing loss of OCM in two mouse strains with contrasting temporal patterns of ARHL, we have demonstrated that OCM and Ca^2+^ buffering play an important role in regulating ARHL. This phenomenon leads us to speculate that Ca^2+^ signaling may be a gateway into a global process that regulates OHC functionality and response to stimuli. The process breaks down during aging leading to silent OHCs (elevated DPOAEs), loss of efferent synapses, OHC loss, and hearing loss. Likely, there are multiple avenues for disrupting this aging process, which leads to the differences in ARHL progression that we see across animal models. By interfering with Ca^2+^ signaling, which is fundamental to many intracellular processes, we were able to expose the process and enhance its destabilization.

## Author Contributions

LC and DS wrote the manuscript. LC, AH, KM, AC, AL and PS performed the tissue preparation, immunostaining and imaging. LC, DS, AH, YY, AC and AL contributed to ABR and DPOAE analysis. All authors discussed and edited the manuscript.

## Acknowledgments

We would like to thank Jack Charles, Jemima McCluskey, Adel Azghadi, and Maria Kazantsev for technical assistance. This research was supported by the following grants to DDS: National Institute on Deafness and Other Communication Disorders (K18 DC013304), a 2015-2016 Fulbright Scholar Award, and an American Hearing Research Foundation grant.

